# Annotating Interchromosomal Interactions at Sub-Megabase Resolution Using Network Clustering Coefficients

**DOI:** 10.64898/2026.01.29.702487

**Authors:** Yingjie Xu, Ian J. Anderson, Rachel P. McCord, Tongye Shen

## Abstract

Specific interchromosomal interactions involve communication between non-homologous chromosomes, enabling coordinated genomic activities such as gene regulation. However, because these communications are often embedded within a nonspecific and noisy background of contact interactions, it is essential to annotate these interaction patterns at the resolution of genomic positions. Such annotation facilitates clean visualization and comparison with linear genomic features to reveal underlying biological functions. We developed and validated a set of network-based metrics as cross-chromosomal interaction descriptors that bridge complex 3D genome structures and 1D functional genomics. By utilizing graph-theoretic representations, these network-based features succinctly summarize complex inter-chromosomal relationships. We constructed a graph representation of contact interactions derived from Hi-C data and implemented three annotations that capture the distinct "many-body" nature of the interactions. Among these, we demonstrate that ΔC4 (a cis-contact-mediated 4-cycle interaction metric) is superior to both ΔC3 (a cis-contact-mediated 3-cycle metric) and C4_E_ (a direct 4-cycle metric of trans contacts) at filtering noise and providing the most straightforward interpretation. Applying these metrics to chromosomes 17, 19, and 22 of the GM12878 cell line, we found clear evidence that different chromosomes rely on a shared set of interaction hot spots to communicate. Overall, this network-based framework reveals distinct chromosomal regulation patches and provides insight into how chromosomes associate with each other and organize relative to the nuclear envelope.

## I. Introduction

The spatial organization of chromosomes within the cell nucleus may affect multiple aspects of gene regulation and determination of cell fate [1–3]. Interactions between chromosomes play an important role in interchromosomal communication and cross-chromosomal regulation of genes [4–7]. Interchromosomal interactions, also known as transchromosomal interactions (trans contacts), are defined as interactions between different chromosomes [5] whereas the more numerous internal contacts are termed cis contact interactions.

Although trans contacts happen far less frequently than cis contacts, they are not random and can contain specific interactions [8–10] which may assist researchers in decoding how chromosomes are spatially organized and how they communicate with each other [5, 11, 12]. Interestingly, imaging techniques reveal that chromosomes from different species at specific points of the cell cycle have distinct contact interaction patterns, ranging from Rabl-like architecture to chromosomal territory [13]. During interphase and within the nucleus, highly regulated chromosomes occupy distinct, localized regions termed chromosome territories (CTs) [14, 15]. These territories reduce the nonspecific interactions between chromosomes, helping preserve the integrity and transcriptional activity of each chromosome [14, 16]. The spatial configuration of these territories both enables and restricts interactions between specific regions across different chromosomes [16–18].

At a higher resolution, interchromosomal interactions can be analyzed using advanced techniques such as Hi-C, a powerful method to explore interchromosomal interactions across the entire genome [19]. Hi-C is a chromosome conformation capture technique that measures the interactions between chromosomal regions within the nucleus at a resolution of up to 1 kb [20]. This method can quantify the frequency of physical association between regions, creating a detailed map of cis and trans contacts [21]. Substantial efforts have been directed towards developing algorithms and computational models to extract information contained in the massive datasets generated by Hi-C and other technologies [22–27]. These computational approaches are essential for identifying key interaction hubs, uncovering functional connections between chromosomal regions, and elucidating the patterns of nuclear organization [28, 29].

By integrating experimental data and computational models, researchers can infer the biological implications of interchromosomal interactions and gain insights into processes such as transcription regulation, DNA replication, and repair [30, 31]. A key area of data analysis involves rendering the 2D Hi-C contact matrix into 1D linear annotations along the genome. One widely adopted existing method is a principal component analysis (PCA)-based A/B compartment analysis, where the sign of the compartment strength (the elements of the top eigenvector) is often associated with the active (A) versus inactive (B) genomic region [21]. Such PCA-based approach primarily focuses on revealing one global feature of cis contacts and the internal organization. However, extending these methods to analyze both intra- and interchromosomal contacts introduces challenges, such as imbalances in cis vs trans contact strength and spotting “few-body” interactions. One can also drastically reduce the resolution and study all interchromosomal contacts using the whole chromosome as a unit, as shown in Ref [15]. However, the specific interactions at gene regulation level are completely lost. Other combined linear and nonlinear metrics may better capture these contact interactions at a fine resolution of megabase or higher. In this work, we examine 3-body and 4-body descriptions that are based on network features.

Abstract graph layouts [32–35] and network analysis [36–38] are powerful mathematical tools for characterizing various complex structures, including biological structures. Here, sophisticated interactions between regions of genome are abstracted as links (edges) between nodes (vertices) [39, 40]. By discretizing spatial connections, quantifying contact interaction strengths, and constructing graph representations of chromosomal structure and interaction, researchers can apply a wide range of network properties to elucidate subtle biophysical and geometrical features contained in genome organization [41]. Both local features (such as degree centrality, eigenvector centrality, local clustering coefficients, [36], and assortativity) and global properties (such as modularity and topological weights) of a network [42] can be used to characterize genome organization.

In this work, we focus on the chromosome structure network (CSN), which designates the linear genome units (such as transcription units) as nodes and the contact interaction between units (such as those indicated by Hi-C data) as “links”, as pointed out by previous studies[40, 43, 44]. Network models of genome structures at other resolutions also exist and each model emphasizes different aspects of genome organization. For instance, in a polymer network model at kilobase resolution, the links symbolize the physical connectivity and topology of dsDNA [42]. In contrast, CSN highlights the noncovalent interaction aspects of the genome organization. A notable application used multigraph representation of interchromosomal interactions in plant cells to demonstrate Rabl-like configuration and how centromeres and distal regions of chromosomes interact [40]. Another study analyzed assortativity and its relationship with specific chromatin states at finer resolution, focusing primarily on interactions within a single chromosome [45]. In this study, we aim to have a quantitative description of the interchromosomal interaction at a resolution of 250 kb using clustering coefficients. Local clustering coefficients (triangle-based clustering coefficient C3 and square clustering coefficient C4) capture many-body interactions by measuring whether three to four nodes of the network form tightly connected clusters of multilateral (trilateral and quadrilateral) interactions.

The earlier network model of chromosomal structure (derived from cis contact only at 50 kb) [46] has already uncovered a set of intriguing properties unseen previously for single chromosomes. As shown in Figure 1, a CSN model of chromosome (based on chromosome 21) contains three parts: a core and two ends. The (network) core contains “urban” nodes that are high in degree centrality, betweenness centrality, closeness centrality, but low in clustering coefficients such as C3 and C4. “Urban” nodes have more neighbors, but they are not connected to each other. In contrast, the “suburban” nodes at the two ends are low in number of neighbors but high in connectivity as shown in higher clustering coefficients. Like a Janus particle, two ends of the structure have different properties. The “stem” end largely contains nodes of inactive genes manifested by high heterochromatin contents while the other end, the “cap”, contains active nodes, low in C4, but high in C3. The stem also contains nodes that are interacting with the nuclear envelope.

**Figure 1.**
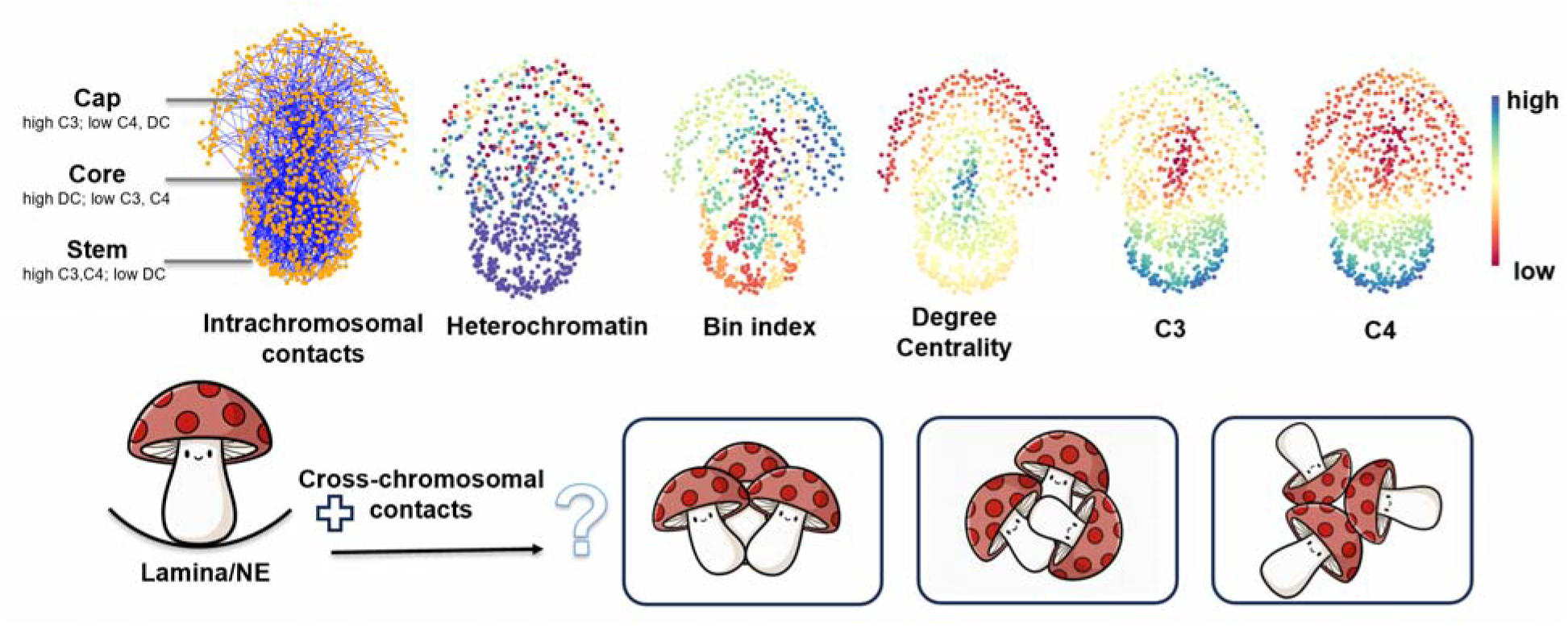
A simple network display of contacts within chromosome 21 (GM12878) Only the Hi-C interaction resolved region (15.25-48.25Mb) is used at the 50-kb resolution (orange dots). Here the top 40% (ranked by strength) of the contacts are shown (down sampled to 1% for a better visualization). Collectively, network information (node-based network centralities and clustering coefficients) provides a picture of structure organization of single chromosomes. With cross-chromosomal contacts being considered, network analysis can assist us to understand the interaction patterns between chromosomes.

The present network model (including trans contact) represents a significant extension of the network approach to capture communication between chromosomes. Our goal is to use node-based network properties to identify key features of contact interactions occurring between different chromosomes. Because the interchromosomal subnetwork forms a bipartite network [37, 47], the C3 metric cannot be directly applied to quantify node-level interaction strength. Instead, we show below that it is possible to develop one direct and two indirect approaches to characterize interchromosomal interactions.

Applying abstract graph and network theory can also bridge the detailed structural features of chromosome territories to specific chromatin state labels [48] and other biophysical annotations such as lamina-associated domains (LADs) [49]. Previously, using a network of cis contacts, we reported that semi-local network properties such as C4 are suitable descriptors that correlate with certain biological features and annotations, such as chromatin states and LADs [46]. The effectiveness of C3 and C4 suggests that biologically relevant interactions can involve three- or four-body contacts, meaning proximity to three or four genomic regions is key for certain biological features. In contrast, metrics such as eigenvector centrality and A/B compartment classifications rely on semi-long or long-range, many-body interactions. While these may provide stronger global indicators of overall structural organization, they can be less responsive to subtle yet biologically meaningful structural variations [50]. When we extend the network to include trans contacts, the model can deepen our understanding of how chromosomes are organized and reveal the underlying patterns that govern interactions between genomic regions of different chromosomes.

In this study, we use a resolution of 250 kilobase (kb), where each genomic bin corresponds to a single node in the chromosome structure network. Because chromosome structures are hierarchical, the corresponding network at one resolution might have structure patterns that are not visible at another [51, 52]. The 250 kb resolution represents a balance between maintaining sufficient contacts for reliable analysis and avoiding excessive averaging over distinct genes—on average, each node spans only a few genes, since human gene density is about 15 genes per megabase.

We demonstrate newly developed metrics for communications via cis contacts between chr17, chr19, and chr22, which has been previously reported to exhibit high frequency of interactions [15, 53]. We limit the analysis solely between pairs of chromosomes and construct each network in a pairwise fashion as shown in Figure 2. This numerical simplification is used in place of directly modeling the multi-chromosomal contact network. However, it is straightforward to generalize this network approach to treat cases where three or more chromosomes form biologically important contact interaction hubs. The pairwise results can serve as a basis for testing how well the shapes of individual chromosomes are defined, and the extent of physical constraint and interaction that promote or inhibit the occurrence of three-body or many-body loci, i.e., simultaneous contacts between three or more chromosomes [54, 55].

**Figure 2:**
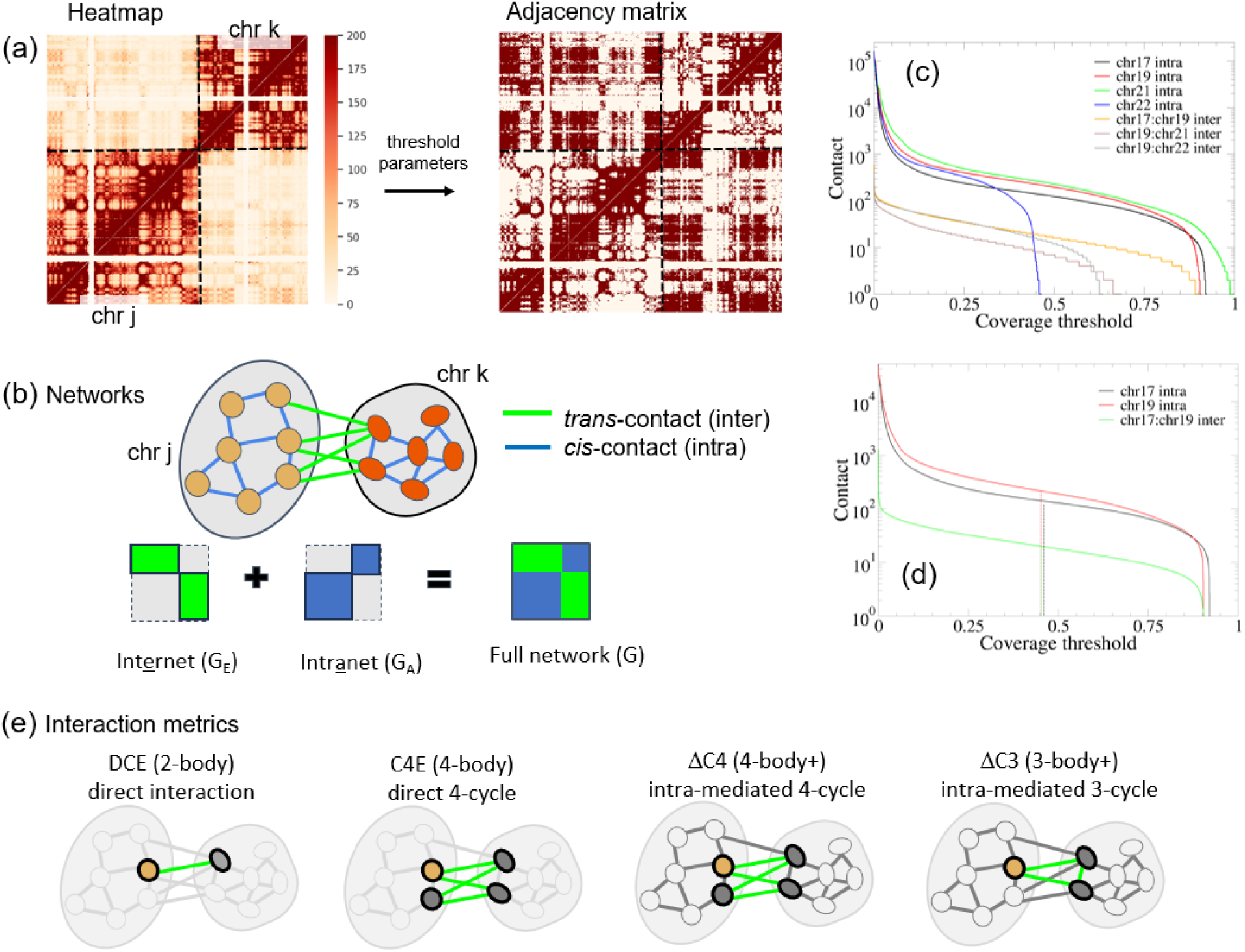
(a) The conversion of contact heatmap of a pair of chromosomes to a chromosome structure network shown. Three individual threshold values are used to ensure internal contact networks of chr j, of chr k, and the interchromosomal network. (b) A cartoon illustration and the corresponding matrix representation of network where the full network can be divided by intra-contact subnetwork (blue, cis-contact links) and inter-contact subnetwork (green, trans-contact links). The green subnetwork is a bipartite network. (c) The ranked contact strength as a function of its ranking (normalized by the size of the contact matrix). Here “cis” indicates the intra-chromosomal contact while “trans” indicates the inter-chromosomal contacts. (d) The corresponding ranked ICE balanced contact value shows the inverse of the distribution function for chr 17 and/or chr 19 systems. The dashed line shows the cutoff position for constructing the inter-chromosomal contact network (the median of all non-zero values). (e) Beyond the direct interaction measured using degree centrality (DC_E_), three interaction metrics (C4_E_, ΔC4 and ΔC3) that filter distinct features of the interaction network are defined.

It also should be noted that because our network is constructed from mean contacts, the direct multi-locus interactions observed in single-cell Hi-C and other related single-molecule methods [20, 56] are not present in this work. Nevertheless, metrics like clustering coefficients still provide valuable insights into the multivalency of contact interactions through a “mean-field” framework based on bulk Hi-C data. Extending beyond the current bulk Hi-C framework, the network analysis on single-cell Hi-C data may provide methods to study the dynamic correlation of contacts.

## II. Method and systems

Abstract graphs represent the essence of structural components of a complex structure and the interactions among components. Generally, such graphs consist of two elements: “node” and “link.” In graph theory, these are referred to as “vertex” and “edge,” while in the physical sciences, they are known as “sites” and “bonds.” In this study of chromosomal structures, each node represents a chromosomal position, which is also called a genomic bin. Each node has a default size of 250 kilobases (kb), unless otherwise stated. The slightly large bin size is chosen as the trans contact is sparse.

Converting from a Hi-C heatmap to a network is equivalent to transforming contact matrix *N_jk_* (where each element represents contact strength between nodes j and k) to adjacency matrix *A_jk_* (where elements are Boolean). The links between nodes denote spatial proximity between chromosomal positions, corresponding to “contact hits” identified in Hi-C experiments. Such link formation can be defined based on a threshold of specific contact strength or frequency measured from Hi-C data, that is, *A_ij_* = 1 if *N_ij_* > *N_c_* and *A_ij_* = 0 otherwise. However, as was argued previously for the cis contact network, selecting a specific percentage of links, network density = 50%, to be formed provides more information for the bulk Hi-C contact matrices. Here, the density of the graph is defined as the number of links divided by the maximum possible number of links in the graph. One reason is that the mean contact matrix of chromosomes is highly resolution-dependent and sensitive to experimental procedures. As a result, it is challenging to establish a consistent and transferable threshold of contact strength when transforming a Hi-C contact map into a chromosome structure network. Instead, we define coverage parameters, which rank relative contact strengths and specify the percentage of links retained from the complete graph.

These coverage parameters provide an intuitive connection to information theory and better reflect the characteristics of Hi-C data. This approach emphasizes the most significant and reproducible contact interactions. It is worth noting that other studies have proposed different criteria for defining “significant” contacts, typically to discern noisy interaction from stable ones [28, 57]. As discussed later in the Results section, our informatic link-percentage criteria are intentionally more stringent in comparison, aiming to minimize numerical noise introduced during network construction.

The link formation is similarly defined as the previous network of cis contacts. Specifically, the threshold of contact interaction strength (after ICE balance [27]) is selected to the median of nonzero elements of the contact matrix. As discussed in the previous work on cis CSN construction [46], this network construction yields a fixed coverage at 50% of maximum possible links, a level that captures the greatest amount of structural information relative to a random network measured by Shannon entropy. For the intr<u>a</u>chromosomal (cis) contact network, G_A,_ a single threshold is used to define link formation. However, for the whole network G, which includes both G_A_ and interchromosomal (trans) contact network, G_E_, the raw “hits” between chromosomes are much weaker (roughly only 10%) than the corresponding item within a single chromosome. Therefore, we employ a separate threshold for each subnetwork.

Using a pairwise system, such as the chr17-chr19 interaction (Hi-C data from ref [20]) as an example, the adjacency matrix of the full network contains three components (cis chr17, cis chr19, and trans chr17-chr19), as illustrated in Figure 2a-b. We have defined the full network G joined by two subnetworks G_E_ and G_A_. Graph G_E_ represents purely direct trans interactions. Because G_E_ excludes all contacts within a single chromosome, it is a bipartite network. In such a network, such as a reader-book or an actor-film network [37, 38, 47], nodes are divided into two groups, and all the links are formed between these groups and not within them.

Bipartite networks do not contain cycles of odd network lengths, resulting in strict zero values for all C3-based (triangle based) clustering coefficients. However, bipartite networks can still be characterized by even-length cycles, such as those quantified by the square clustering coefficient (C4). To incorporate triangle-based clustering information, we construct a reference graph G_A_, containing only internal (intrachromosomal) interactions, and a full graph G = G_E_ ∪ G_A_. Both G_A_ and G yield nonzero C3 values. The difference between them presents the effect of G_E_ measured by triangle-based clustering coefficients. Finally, we denote the clustering coefficient of the full network G as C3, that of G_A_ as C3_A_, and that of G_E_ as C3_E_. Similarly, the C4 values for G, GA, and G_E_ are denoted as C4, C4_A_, and C4_E_, respectively. We can then define ΔC3 = C3 − C3_A_, representing an alternative measure of contact interaction. Additionally, we can also calculate ΔC4 = C4 − C4_A_. As illustrated in Figure 2e, three node-based metrics, C4_E_, ΔC3, and ΔC4, were used to evaluate contact strength at the genomic-bin resolution. Among these, C4_E_ directly quantifies interchromosomal contacts, whereas ΔC3 and ΔC4 capture cis-contact–mediated trans interactions.

The rationale for studying chromosomes 17 and 19 is that they exhibit strong interchromosomal interactions. In GM12878 cells, chr17 exhibits the strongest trans contact interactions, followed by chr19. The cis interaction between chr17 and chr19 is stronger than any other pair of chromosomes. Therefore, our analysis centers on the interaction network formed between these two chromosomes. Previously, the overall contact PCA results have also demonstrated that the dominant eigenvector (PC1) has large loading coefficients from chromosomes 17, 19, 22, and 1 [15].

In this study, we examine several key linear biological annotations including lamina-associated domains (LADs) [58] and chromatin states [59]. LADs are large chromatin regions anchored to the nuclear lamina, typically corresponding to transcriptionally repressed zones that influence chromosomal positioning and functional organization within the nucleus. Chromatin states, defined by distinct patterns of histone modifications, mark genomic regions of active or repressed gene expression and provide insight into regulatory mechanisms across the genome. Understanding the connection between these annotations and trans interaction strength is crucial for unraveling the complex network of chromosomal interactions that maintain cellular function and genomic integrity.

## III. Results and discussion

### 1. Construction of interaction networks using Hi-C contact maps

The data of this study came from the Hi-C sequencing dataset from human lymphoblastoid cell line (GM12878) reported by Rao et al. (2014) at 250-kb resolution [20], unless otherwise specified.

As described in the methodology, two main factors influence link selection: (1) each contact must be statistically meaningful (i.e., exceed a significance threshold to avoid random “hits”), and (2) when a large fraction of contacts is meaningful, selecting approximately 50% of them minimizes information loss during the transformation from the continuous contact matrix to a discrete adjacency matrix. In this setup (chr17–chr19, 250 kb bin size), we find that a large fraction of elements in the contact matrix contain nonzero values. We first examine a ranking contact function *f(x)*, where *x* represents the normalized rank index (rank index divided by the total number of contact pairs, sorted from largest to smallest of contact strengths), and *f* returns the corresponding contact strength (*n_ij_*, the number of raw contact hits). Mathematically, this ranking function corresponds to the inverse of the cumulative distribution function F(*n_ij_*), and its derivative yields the probability density of p(*n_ij_*).

As shown in Figure 2a, ranking contact functions for seven networks are shown: intrachromosomal contact matrices for chr17, chr19, chr21, chr22 and interchromosomal contacts for chr17-chr19, chr17-chr22 and chr19-chr22. For the example of the chr17-chr19 system, the linear sizes of the chromosomes are N_17_ = 325 and N_19_ =237. Thus, the total number for intrachromosomal contact for chr17 is N_17_× (N_17_ −1)/2 = 325×324/2= 52,650 whereas that for chr19 is N_19_ × (N_19_ -1)/2=237×236/2 = 27,966. The total interchromosomal contact pairs N = N_17_× N_19_ = 77,025. For the cis contacts of chr17, the ranking function drops to zero at x= 0.923, indicating a maximum coverage of 92.3%. At this maximum coverage, even a single Hi-C hit is sufficient to form a link. The corresponding values for internal chr19 and chr17-chr19 are 91.5% and 90.4%, respectively. Both interchromosomal and intrachromosomal contacts exhibit roughly 90% maximum coverage. However, cis and trans contact strengths are quite different. The percentages of strength (represented by total raw contact hits) for cis chr17 contacts, cis chr19 contacts, and trans chr17-chr19 contacts are 55.7%, 42.1%, and 2.2%, respectively. The ratio of intra- vs inter-contact strength is approximately 50:1 for this specific sequencing data. Similar ranking curves can be plotted for chr19 and chr17–chr19. As observed, interchromosomal contact coverage is lower than intrachromosomal coverage, evident from the faster decay of its ranking curve toward zero. Instead of using the raw *n*□□ data directly, we apply ICE normalization [27], which transforms the raw contact counts *n*□□ into balanced values *a*□□. As shown in Figure 2b, the contact strength *a*□□ for the chr17–chr19 system is plotted against the normalized rank index. The cutoff threshold is defined as the median of all nonzero contact values. To ensure that our observations are generalizable, we also studied chr22 (N□□ = 205) and its interactions with chr17 and chr19.

### 2. Assessment of network metrics for interchromosomal interaction strength

As described in the Methods section, we constructed three node-based metrics (C4_E_, ΔC3, and ΔC4) designed to quantify the strength of interchromosomal interactions. As mentioned in Methods, C3E cannot be used since G_E_ is bipartite. We first evaluated if these three metrics identify the consistent features of interchromosomal interactions (Figure 3).

**Figure 3:**
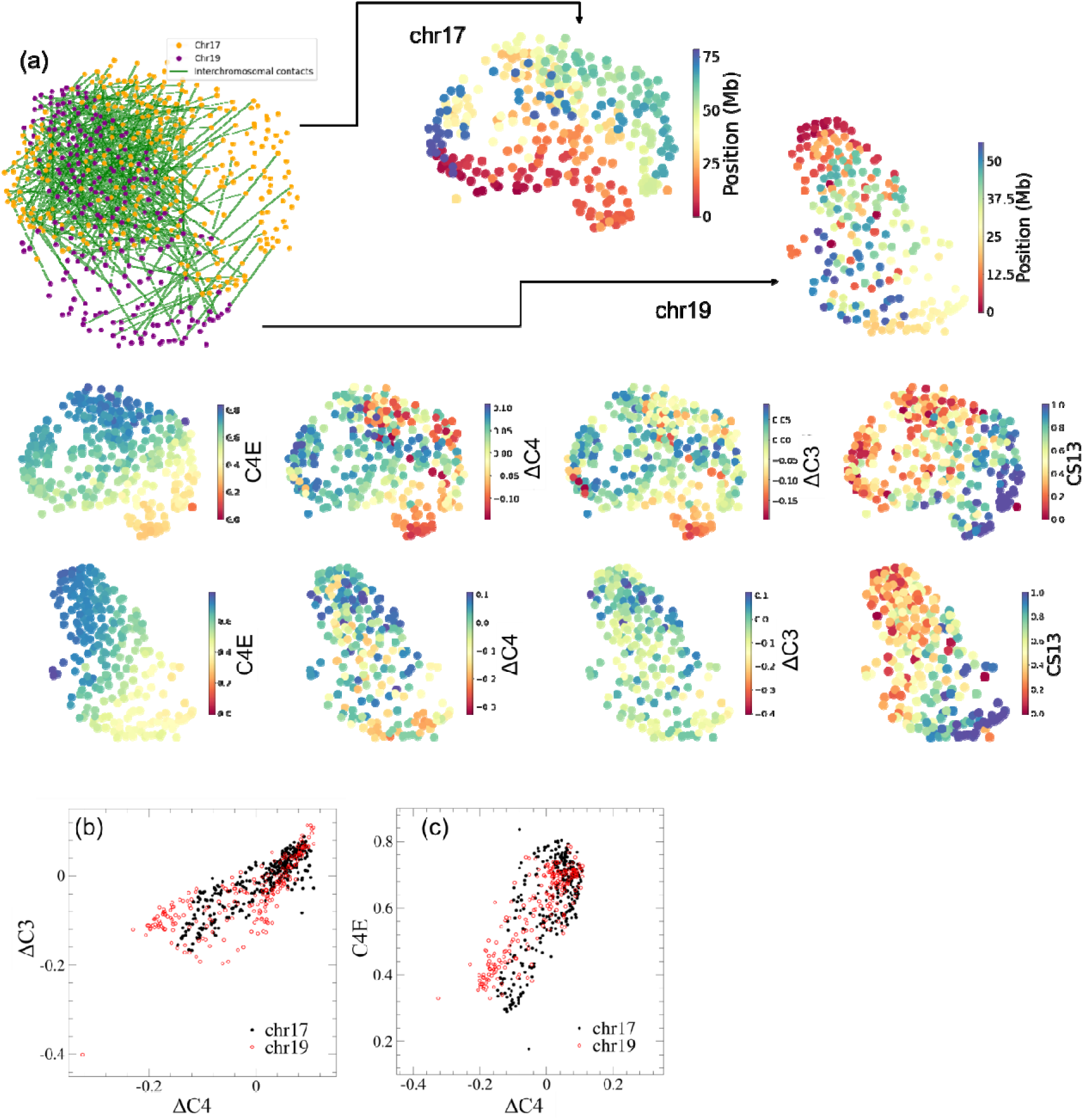
(a) The top left image shows the force distribution layout of nodes in chromosomes 17-19 system, and the other panels show two chromosomes separately. A randomly sampled 1% of the top 50% interactions (edges) are shown to improve visual clarity. The rightward arrows point to chromosomes 17 and 19 with nodes colored by their genomic positions. Below, from left to right, nodes are colored by C4_E_, ΔC4, ΔC3, and the intensity value of the chromatin state CS13 signal (heterochromatin). (b) The scatter plot of ΔC4 versus ΔC3 of the same Chr17:Chr19 system. (c) The corresponding scatter plot of ΔC4 versus C4_E_.

All three metrics for chr17-chr19 are visualized on a force-directed network layout in Figure 3a. Additionally, these metrics plotted as functions of the genomic bin are shown in Supplementary Information (SI) Figure S1. Further, additional results for the corresponding chr17-chr22 and chr19-chr22 systems are provided in SI Figure S2. The network layouts were generated using the Kamada–Kawai algorithm [32] The genomic bin indices were plotted alongside the network to serve as reference coordinates. The strongest interchromosomal communication originates from the p-arm of chr19 and the distal regions of both the p- and q-arms of chr17. As shown in Figure 3a, nodes with high metric values coincide with regions exhibiting dense interchromosomal interactions, corresponding to the top 50% of contact strength. For clarity, not all top 50% of interactions are plotted; instead, 1% of these links were randomly selected and are shown explicitly as green lines. When only the top 1% of interactions are considered, there is less correlation between metrics such as C4_E_ and link density (SI Figure S3). This observation suggests that the strongest direct interchromosomal contacts rarely participate in extensive multivalent interactions. Scatter plots (Figures 3b and 3c) illustrate the relationships among three interaction metrics: C4_E_, ΔC3, and ΔC4. The results reveal positive correlations among the three metrics. Therefore, all three network properties are largely consistent and measure the inter-chromosomal interaction strength in certain aspects. Deviations from a linear model among these metrics may reflect specific metric emphasis and/or structural features of a specific genome, as further demonstrated in Figures 4 and 5.

**Figure 4:**
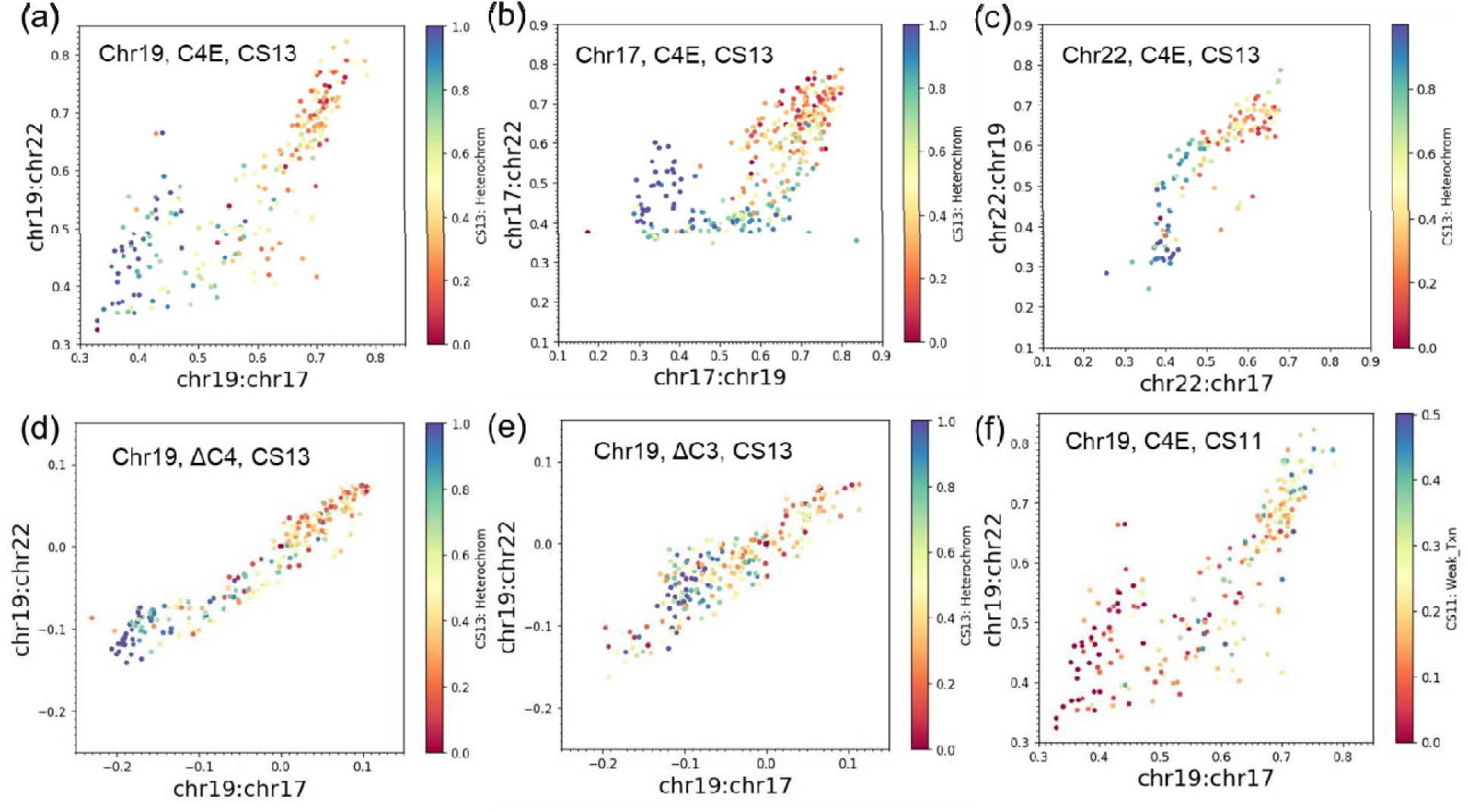
Scatter plots of communication metrics of a specific chromosome colored by chromatin state annotation. (a) The scatter plot shows how chr19 interacts with the two other chromosomes (x-axis with chr17, y-axis with chr22), expressed by C4_E_ and color coded by the chromatin state CS13 (intensity of the Heterochromatin signal of the fragment). Each node represents a 250kb genomic bin from chr19. Similarly, we display other situations with a combination of different chromosome, communication metric, and chromatin state in (b–f).

**Figure 5.**
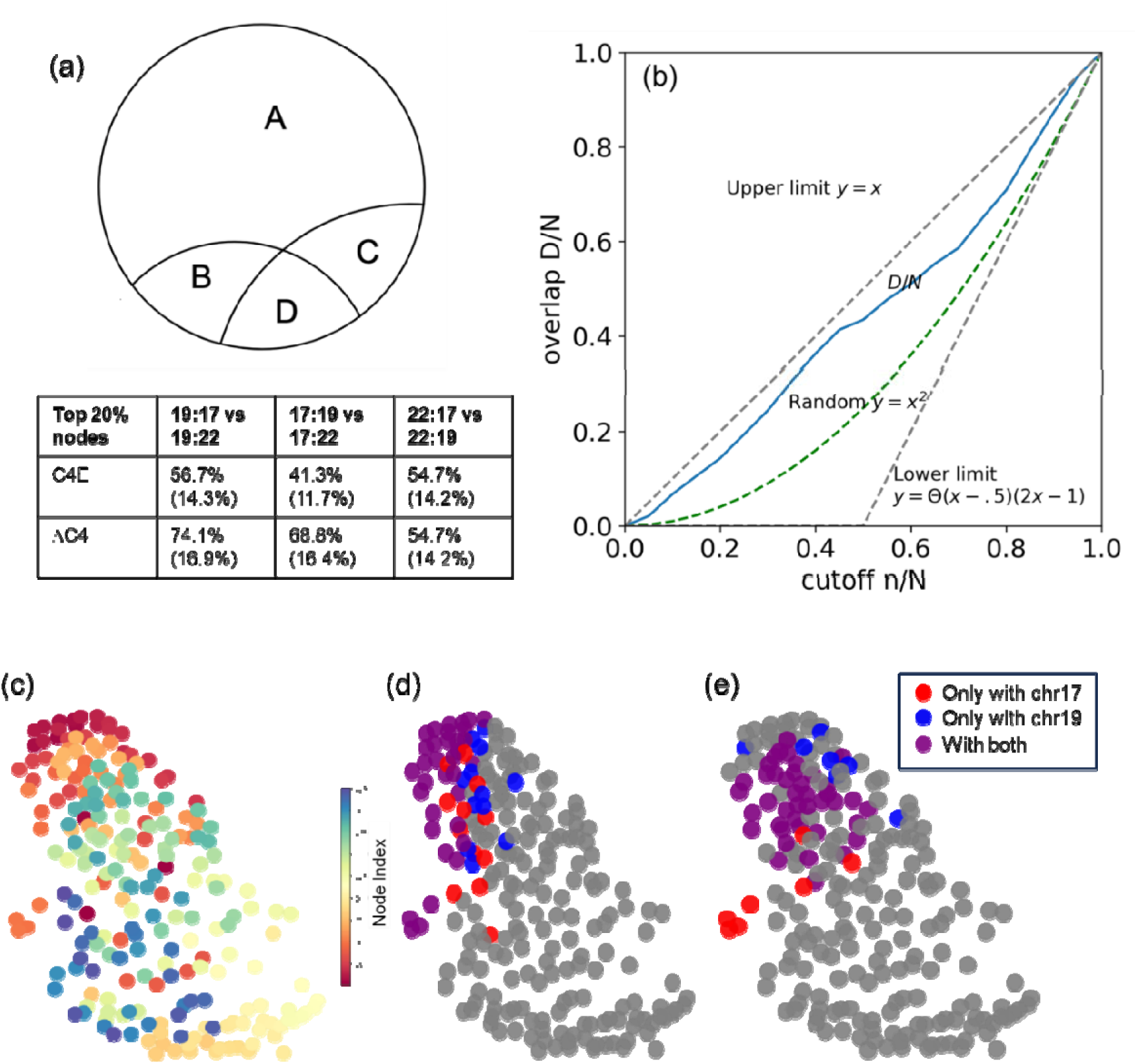
(a) Concurrence parameter shows the percentage of the chromosome nodes that are simultaneously involved in interactions (judged by node-based metrics) with two other chromosomes. Here A indicates a subset of nodes that does not interact with either of two other chromosomes. B and C indicate sets that interact with only one other chromosome. D indicates a set that interacts with both. We define concurrence parameter = D/(B +C +D) = D/(CD + BD – D). The top 20% values of C4_E_ or ΔC4 were selected as the cutoff value for contact. The percentages in brackets indicate the proportion of the D region to the entire chromosome. (b) The normalized overlap as a function of normalized cutoff. (c) Nodes of chromosome 19 isolated from the chromosomes 17-19 layout colored by bin index. (d) Nodes of chromosome 19 that interact with chromosomes 17 and 22 through different regions. Red indicates contact only with chr17. Blue indicates contact only with chr22. Purple indicates contact with both. The top 20% values of C4_E_ were selected as the cutoff value for contact. (e) Same setup as panel (d) except ΔC4 is used as the interaction strength.

For these constructed CSNs, it is also interesting to explore the relationship between other node-based properties, such as the clustering coefficient (C3) and degree centrality (DC). For example, a hallmark of certain biological networks—including protein-protein interaction, metabolic, and gene regulatory networks—is a negative correlation between C3 and DC of nodes [60–62]. This fingerprint indicates that certain networks possess a hierarchical and modular structure, even under the refinement of degree correlation [63]. In contrast, nonhierarchical networks (such as technical networks) might be flat or even exhibit positive correlation between C3 and DC at the low-to-mid range of DC. It is possible to have a multiscale graph with positive correlation at low DC values and negative correlations at large scales (high DC) involving “global hubs” which resulted in a “chevron-shaped” correlation. Hybrid and multiscale networks were proposed for systems such as certain protein interactions and semantic web among others [64–66].

We found that such a hybrid network fits the description of the cis network (G_A_). As illustrated in Figure 1, when we ranked the three regions of the network (cap, stem, and core) by DC values, we found DC_cap_ < DC_stem_ < DC_core_. In contrast, when examining C3, we found C3_core_ < C3_cap_ < C3_stem_ for CSN_A_. Thus, at the low-mid range of DC values (comparing cap and stem), a larger DC is related to a larger C3, ensuring a positive relationship. Conversely, at the high-DC end (comparing stem and core), a negative correlation is shown. Since the trans network (G_E_) does not have non-zero C3E, we compared instead C4_E_. Again, as shown in SI Figure S3b, complex patterns are shown between C4_E_ and DC for G_E_.

### 3. Metric ΔC4 is superior for describing the hubs for interchromosomal interactions

After defining the metrics that represent the strength of cis interactions, we would like to compare these descriptors and address one important question regarding cross-chromosomal interaction: does a given chromosome use the same genomic region(s) to interact with other chromosomes? To explore this, we generated scatter plots to examine whether a chromosome, such as chromosome 19, uses a set of shared nodes to interact with other chromosomes, including chr17 and chr22, as shown in Figure 4. In addition to plotting the interaction strength, we incorporated a third variable, chromatin-state appearance, by color-coding.

The chromatin state appearance, denoted Q_j_, for the *j*-th nodes is defined as follows. We begin with labels of chromatin states assigned by Ernst et al. [59] for GM12878 cells, where each position (resolution = 200 bp) of chromosome is assigned a single state CSX (X = 1 to 15). To quantify how frequently represented a specific chromatin state is, we assign the *j*-th node the chromatin state appearance for X-th chromatin state, CSX, as Q_j,X_ which is normalized to a value between 0 and 1. Thus, appearance Q of a node is defined as the ratio of the total sequence length where a CS label is detected in a node to the total length of the node (L =250 kb), Q *= L_X_ / L*. If none of the specified CS intervals fall within the node, Q is strictly zero. We further denote the total number of nodes with Q = 0 as N_o_. If N_o_ > 90% × N_19_, we deem the specific CS state does not have enough representation in chr19, and we omit such a CS from further analysis to avoid spurious correlation between chromatin state labels and network features. For chr19 of cell line GM12878, we found that CS13 (heterochromatin) has the highest presence among 15 CSs (94.9% of the nodes have non-zero values for CS13), followed by CS8 (86.9%) and CS11 (86.5%), whereas CS3 displays the lowest representation, 35.0% (SI Table S1). Thus, for this cutoff of 90%, all 15 CSs have sufficient hits for chr19 at 250 kb resolution.

From the scatter plot, the interactions (measured by C4_E_) of chr19:chr17 and chr19:chr22 generally show a positive correlation. The nodes clustered on the diagonal line represent genomic bins on chr19 that interact similarly with both chr17 and chr22. However, certain off-diagonal features can be observed in the graph. The bins clustered to the upper left of the diagonal line show a stronger tendency to form contact with chr22 and less so with chr17. We observe that the nodes on chr19 that frequently contact chr17 are likely transcriptionally active according to the low CS13 appearance (red color nodes) on Figure 4a. Similar patterns are observed for chr17 and chr22, as shown in Figure 4b and 4c, respectively. When we use ΔC4 instead of C4_E_, we also obtained consistent results, as shown in panels 4d and 4e, respectively. Notably, ΔC4 exhibits stronger correlations than C4_E_. Additionally, ΔC4 differentiates CS13 better than ΔC3, which lacks a defined color scheme for the scatter plot. We also show a few other chromatin states, such as CS11 (weak transcription), as shown in Figure 4f. The pattern of CS11 shows an opposite trend since CS11 labels active regions.

From the striking linear relationship in Figure 4d, we deem that ΔC4 is the superior descriptor than ΔC3 and C4_E_, since it demonstrates the most linear behavior between how the same node of one chromosome interacts with two other chromosomes and additionally, it also has the clearest separation between heterochromatic and euchromatic states by color labels. The reason behind ΔC4’s success is that the 4-body interaction descriptor is more robust than 2-body descriptors (such as degree centrality that only counts neighbors) at filtering generic contacts and background noise. While direct 3-cycles yield zero for bipartite structures, the inclusion of indirect, mediated interactions in ΔC4 effectively captures intra-interaction signals that C4_E_ misses.

The reason that ΔC4 also outperforms ΔC3 is the following. While the full network *G* does accumulate new mixed triangles where a single locus on one chromosome bridges an internal cis-contact on another, the 3-cycle geometry is inherently limited by a 1-to-2 node asymmetry. It fails to reflect mutual, localized cluster-to-cluster interfaces. In contrast, a 4-cycle allows a balanced 2-to-2 geometry (Figure 2e) where a structural neighborhood on Chromosome j cooperatively interacts with a structural neighborhood on Chromosome k. Consequently, ΔC4 provides a significantly cleaner, symmetric signature of true interchromosomal hot spots, whereas ΔC3 remains heavily distorted by local cis-clustering noise (of Chromosome k).

To quantify the extent to which chromosomal regions are simultaneously engaged in interactions with other chromosomes, we define a concurrence parameter (Figure 5a). Each node was categorized according to its interaction partners as follows: A – nodes that do not strongly interact with either of the two other chromosomes; B, C – nodes that interact with only one other chromosome; D – nodes that strongly interact with both chromosomes B and C. The concurrence parameter was computed as Concurrence = D /(B +C +D), which represents the proportion of inter-chromosomal interactions that occur concurrently with two partners. Equivalently, the parameter can also be expressed as Concurrence = D /(BD +CD − D). Taking chromosome 19 as an example, BD is the total number of bins interacting with chromosome 17, CD is the number of bins interacting with chromosome 22, and D is the subset of regions involved in both interactions. Only the top 20% of strongly interacting nodes (ranked by network properties) were included in the analysis to reduce background noise.

As shown in Figure 5, when filtering based on C4_E_ values, 56.7% of the regions on chromosome 19 that interact with either chromosome 17 or chromosome 22 are concurrently in contact with both. These regions represent 14.3% of the entire chromosome 19. Figure 5b illustrates how these conclusions vary depending on the selected cutoff thresholds. The upper and lower bounds, and the uncorrelated reference plots are also shown. Figure panels 5c–e depict a distinctive spatial organization pattern of chromosome 19 in its interactions with chromosomes 17 and 22. Specifically, several nodes near the telomeric region of the p-arm on chromosome 19 appear to contact both chromosomes 17 and 22, suggesting the presence of shared contact domains that may function as hubs for interchromosomal communication. Overall, Figure 5 demonstrates different chromosomes (in this case, 17 and 22) share a consistent set of hotspots (local interact environment) for chr19. Our data shows overlaps exceeding 50% (reaching 70% in some cases) when we use ΔC4 as the metric. These shared regions form large, consistent patches, whereas the differences appear more scattered and likely attributable to noise.

Extending beyond the current pairwise framework, multi-chromosomal interactions could potentially be detected through a clustering coefficient of three chromosomes. For example, one can compare C3E_h_ of chr19, which records heterogeneous triplet between chr17, chr19, and chr22 of the networks G and that of a corresponding random network G*[38] and inform us if shared nodes of chr19 makes chr17 and chr22 indirectly attracts (C3E_h_ > C3E_h_*) or repels at these positions. In addition to these overlapping domains, our results also identify distinct regions on chromosome 19 that interact exclusively with either chromosome 17 or 22. Such exclusivity suggests a spatial and functional partitioning within chromosome 19, where some nodes maintain selective contact while others participate in shared interchromosomal interactions.

Furthermore, we quantified the connection between interchromosomal interaction strength (expressed by a linear fit model of ΔC4 and C4_E_) and the appearance of CS labels Q using a correlation coefficient r. The correlation coefficient r, bounded between –1 and 1, indicates the degree to which interaction strength (e.g., ΔC4) is correlated with specific chromatin states (CSs). We first performed a principal component analysis (PCA) on the scatter plot of ΔC4 versus C4_E_, where the first principal component (PC1) defines the direction of consensus between the two node-based metrics. Next, we calculated Spearman’s rank correlation coefficients between the PC1 projections and the chromatin-state appearance values Q. Using chromosome 19 as a test case, we found that chromatin states associated with strong transcriptional activity, such as active promoters (CS1) and strong enhancers (CS4 and CS5), exhibited the highest positive correlations, which indicate these 4-body descriptors resonate with the many-body nature of the corresponding biological processes. In contrast, weaker regulatory states like poised promoters (CS3) showed lower, though still positive, correlations. Conversely, heterochromatin (CS13) was the only chromatin state showing a negative correlation with these metrics (see SI Table S1).

### 4. Annotation of LAD and the association of chromosome and nuclear lamina

In addition to chromatin state labels, lamina-associated domain (LAD) represents another key binary classification. LAD labels where chromosomes are associated with lamina (skeletal and anchoring proteins of the nuclear envelope) [49, 67]. Because lamina association imposes mechanical constraints and defines nucleus morphology, we compare how LAD labels are correlated with network metrics. Our previous study of LADs and cis contacts [46] has already revealed that LADs are located at nodes with high C4A values and C4A is a good feature for LAD classification. Here we would like to explore the connection between LAD and trans contacts.

As the LAD classifier provides binary information for each node (either associated with the nuclear lamina or not), we use cumulative distribution function (CDF) to evaluate the correlation. As shown in Figure 6, three sets of network node-based properties are evaluated to examine how well they are connected to the LAD labels. In each panel, a large separation between the CDF plots (black versus red) indicates the metric shown on the x-axis is a good LAD classifier. Specifically, we compared C4_A_ (for Chr21, a re-plot of the results of Reference [46]), C4_E_, and ΔC4. Among them, LAD status is associated with lower C4_E_ but higher C4_A_. Another inter-chromosomal strength indicator, ΔC4, shows the same trend as C4_E_. These findings suggest that chromosomal regions engaged in interchromosomal contacts are less likely to be located at the nuclear lamina. In contrast, previous analysis of intrachromosomal contact networks using C4_A_ revealed that highly connected nodes are more frequently associated with LADs.

**Figure 6:**
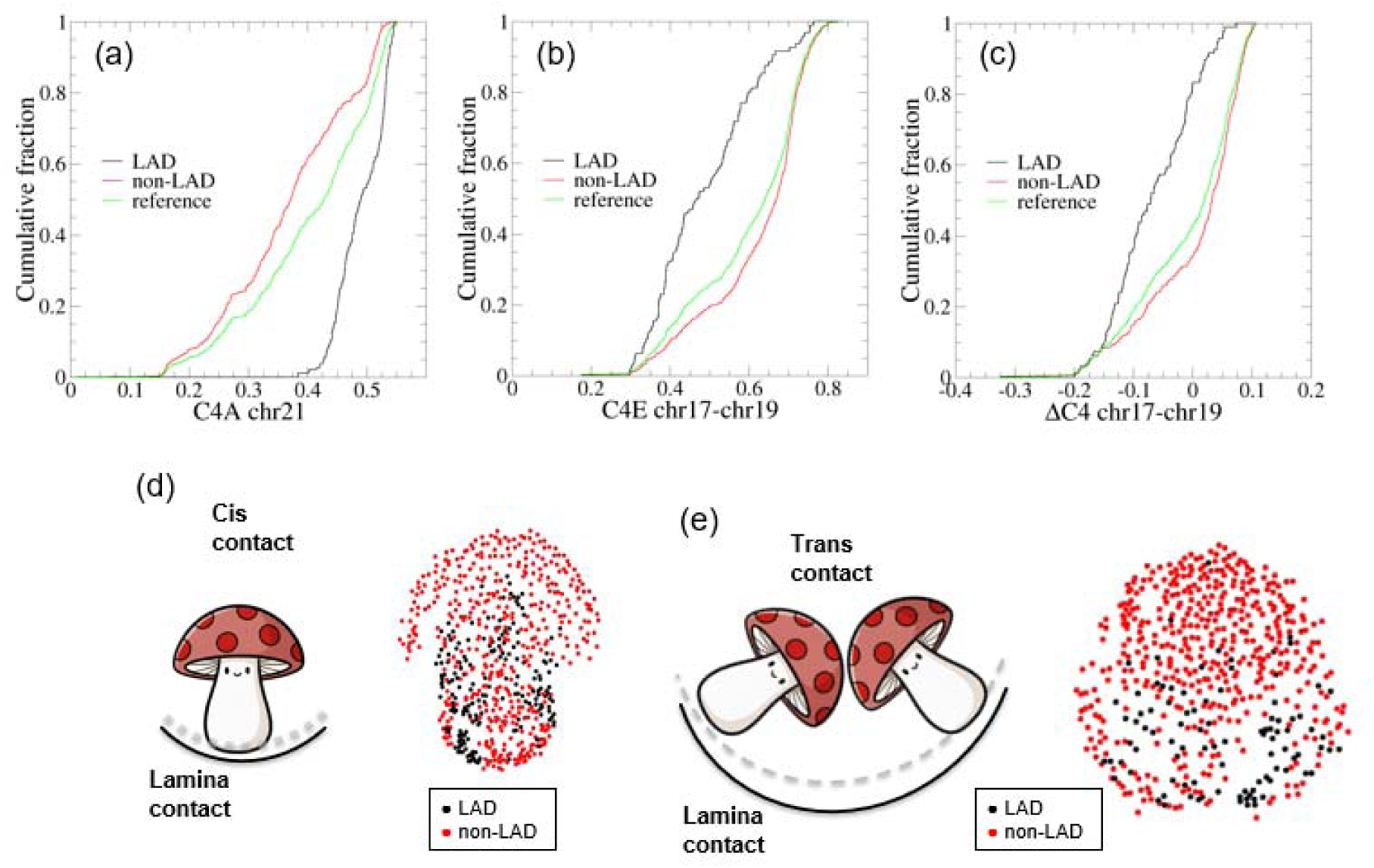
Cumulative distribution fraction (CDF) plots of network metrics for chromosome networks. Here classifier performance concepts using CFP (black), CFN (red), CFR (green) are shown for chr21 using feature C4A in (a), for chr17-chr19 using C4_E_ in (b), for chr17-chr19 using ΔC4 in (c). (d) A mushroom diagram illustrating the interaction patterns of the single-chromosome system (chr21) with the nuclear envelope and the corresponding distribution of lamina-associated domains (LADs). (e) A mushroom diagram illustrating the interaction patterns of the two-chromosome system (chr17-chr19) with the nuclear envelope and the corresponding distribution of LADs.

### 5. Effects of condensin II deletion

Equipped with these three metrics, we further investigated how condensin II deletion mutant (ΔCAPH2) of HAP1 cells influences interchromosomal interactions. Condensin II is known for its importance in chromosome looping and compaction [68, 69]. The absence of CAP-H2 results in extensive changes to chromosome structure and notably affects interactions between chromosomes [70]. The previous intra-chromosomal network analysis reveals increased fluctuation of C3 and C4 as functions of bin index (as shown in SI Figure S4b and SI of reference [46]) in mutant cells, indicating region-specific impacts of condensin II loss on chromosomal connectivity, consistent with the disruption of condensin-mediated loops. In the current study, we extended the analysis to trans interactions. We constructed the chr17–chr19 interaction network using structural data from both wild-type and ΔCAPH2 mutant HAP1 cells during mitosis. The comparison of network properties is shown in SI Figure S4, and the network display is presented in SI Figure S5.

Our analysis indicated that metrics ΔC3 and ΔC4 do not show substantial overall increases or decreases, and the general features persisted across conditions (SI Figure S4c–e). However, in mutant cells, the structural features, represented by peaks and troughs, were noticeably dampened relative to wild-type cells; peak values were reduced, troughs elevated, and the overall curve appeared flattened. These results suggest that mutant cells exhibit a more homogenized chromosomal interaction pattern.

Compared to the internal changes, the changes between chromosomes presented here are less pronounced. Nevertheless, the result is consistent with the known function of condensin II, which primarily affects the ability of individual chromosomes to fold and organize themselves into compact territories. While its role in inter-chromosomal interactions is relatively smaller, the effects are still noticeable: mutant ΔCAPH2 exhibits reduced heterogeneity of inter-chromosomal interaction compared to that of HAP1 cells. These findings indicate that condensin II contributes to the maintenance of a dynamic and heterogeneous nuclear architecture. Such a role is consistent with the reported function of condensin II in human cells [71, 72]. Notably, for the region near the centromere of chr17 (around 25Mb), a peak appeared in the mutant and indicated a possible change of interaction near the centromere. These findings further support the conclusion that condensin II both affects trans and cis interactions [73–75].

## IV. Concluding remarks

Our work focuses on the development of a set network features, (C4_E_, ΔC3, and ΔC4), as quantitative descriptors for annotating specific trans contact interactions between chromosomes and its validation using heterochromatin states and LADs. We found that interaction strengths, characterized by network metric ΔC4, can provide a unified description on how chromosome 19 interacts with two other chromosomes, 17 and 22. Such metrics can also exhibit a definitive association with heterochromatin state as well. In addition, such interaction descriptions can be compared with the annotation for nuclear lamina affinity to gain insights on the spatial chromatin organization, where our application suggests chromosome–nuclear periphery interactions suppress interchromosomal contacts, Using the “produce” diagram analogy, “stems” interact with nuclear lamina whereas “caps” interact with each other. When these interaction metrics are applied to gain insight of mutation effect on genome structure using the ΔCAPH2 mutant as an example, attenuated specific trans-interactions were observed compared to that of haploid HAP1 cells, which corroborates with the previous report that a reduced internal structural heterogeneity due to the mutation.

Overall, this study establishes a network-based framework for annotating interactions between chromosomes using clustering coefficients, offering new insights into chromatin states and chromosome-nuclear lamina interaction through the lens of interchromosomal transitivity. The methodologies and findings provide a foundation for future investigations into nuclear architecture and its implications in development, differentiation, and disease-related chromosome reorganization.

## Acknowledgements

We acknowledge insightful discussions with P. Das and D. Foutch. This research was supported in part by the National Institute of General Medical Science grant R35GM133557 to RPM.

Supplementary Material includes one table and seven figures. Table S1 lists the correlation strength between trans-interaction strength and chromatin states. Figure S1 shows network properties of subnetworks. Figure S2 shows graph representations of additional chromosome systems. Figure S3 shows network graphs using top-ranked links. Figure S4 shows HAP1 WT versus ΔCAP-H2 network property comparisons. Figure S5 shows HAP1 WT versus ΔCAP-H2 network graphs. Additionally, example python notebook codes for network visualization and clustering coefficient calculation can be found on GitHub (https://github.com/tshen-camp/CSN).

