## Supplemental File for "Annotating Interchromosomal Interactions at Sub-Megabase Resolution Using Network Clustering Coefficients"

Supplementary Information for *Annotating Interchromosomal Interactions at Sub-Megabase Resolution Using Network Clustering Coefficients*

Yingjie Xu^1^, Ian J. Anderson^2^, Rachel P. McCord^3^, and Tongye Shen^3*^

1 Genome Science and Technology, Bredesen Center, The University of Tennessee Knoxville, 2 Pratt School of Engineering, Duke University, 3 Department of Biochemistry and Cellular and Molecular Biology, The University of Tennessee Knoxville

Table S1: Chromatin state coverage and correlation with interaction strength on chromosome 19. For each chromatin state (CS1–CS15), the table shows the fraction of bins with non-zero appearance and the Spearman correlation (ρ) between Q and the consensus of communication metrics (first principal component PC1 of ΔC4 vs. C4_E_).

| Chr19 Chromatin State | | Appearance Q (%) | ρ (ΔC4) | p-values | ρ (C4E) | p-values |
| --- | --- | --- | --- | --- | --- | --- |
| CS_1 | Active promoter | 79.3 | 0.601 | 1.18E-24 | 0.782 | 1.89E-45 |
| CS_2 | Weak promoter | 83.5 | 0.577 | 1.96E-22 | 0.709 | 7.99E-33 |
| CS_3 | Inactive/poised promoter | 35.0 | 0.420 | 1.55E-11 | 0.382 | 6.79E-11 |
| CS_4 | Strong enhancer | 63.7 | 0.670 | 3.86E-32 | 0.746 | 1.26E-44 |
| CS_5 | Strong enhancer | 64.1 | 0.614 | 6.02E-26 | 0.672 | 1.68E-33 |
| CS_6 | Weak/poised enhancer | 85.2 | 0.542 | 2.44E-30 | 0.776 | 6.07E-45 |
| CS_7 | Weak/poised enhancer | 83.1 | 0.623 | 7.18E-27 | 0.707 | 2.20E-37 |
| CS_8 | Insulator | 86.9 | 0.486 | 1.84E-15 | 0.476 | 8.39E-14 |
| CS_9 | Transcriptional transition | 62.9 | 0.590 | 1.26E-23 | 0.613 | 4.34E-27 |
| CS_10 | Transcriptional elongation | 76.8 | 0.388 | 6.08E-10 | 0.550 | 3.20E-18 |
| CS_11 | Weak transcribed | 86.5 | 0.420 | 1.44E-11 | 0.672 | 8.14E-29 |
| CS_12 | Polycomb repressed | 51.9 | 0.455 | 1.71E-13 | 0.384 | 2.40E-10 |
| CS_13 | heterochrom | 94.9 | -0.519 | 1.01E-17 | -0.422 | 1.29E-13 |
| CS_14 | Rep/CNV | 56.5 | 0.007 | 0.912537 | 0.169 | 0.024336 |
| CS_15 | Rep/CNV | 39.2 | 0.108 | 0.098681 | 0.165 | 0.022638 |

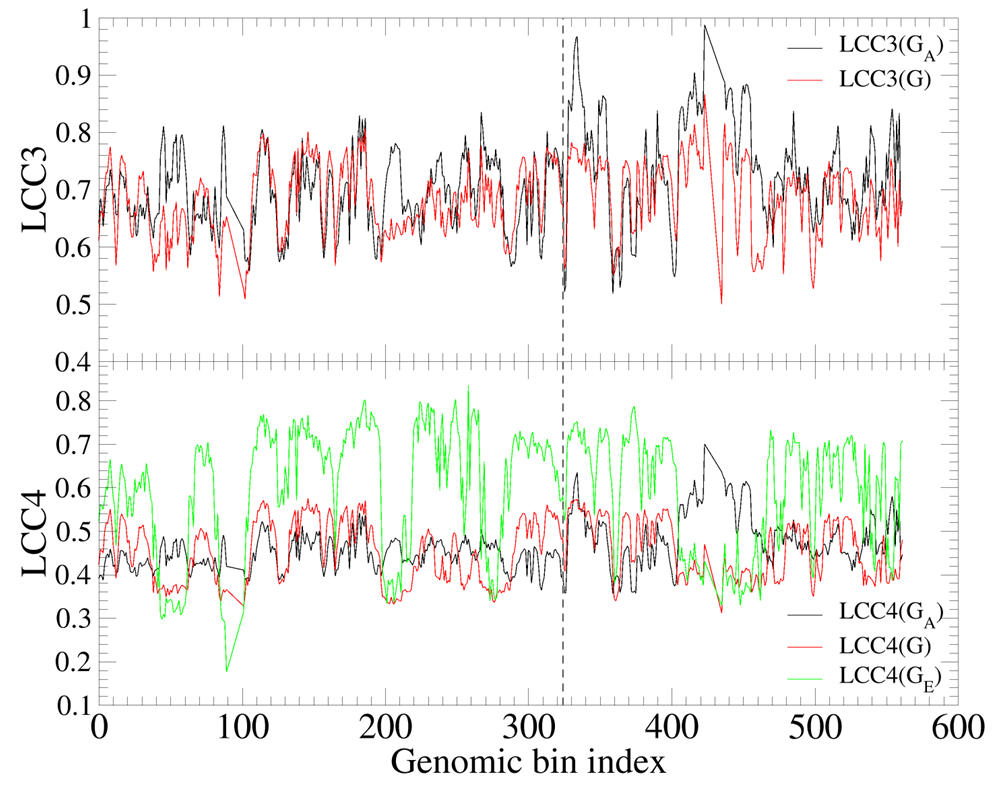

Figure S1: Comparison of LCC3 (C3) and LCC4 (C4) values of different contact networks in Chr17-Chr19 system. Bin indices 0-324 correspond to chromosome 17 (0-81,195,210 bp), and indices 325-561 correspond to chromosome 19 (0-59,128,983 bp), at a resolution of 250 kb.

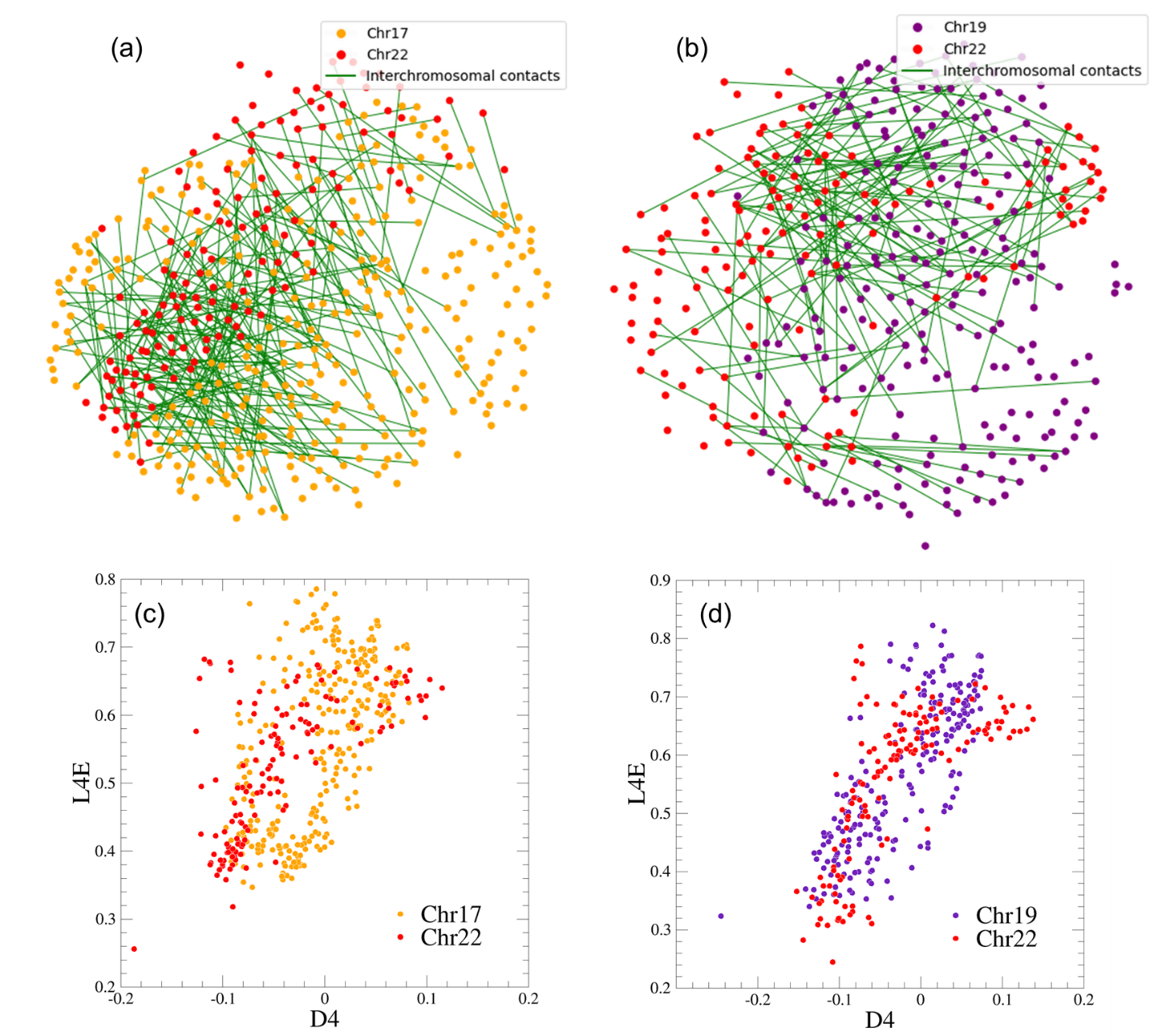

Figure S2: (a-b) Force distribution layout of the (a) chr17-chr22 and (b) chr19-chr22 contact network system in HAP1 wild-type and ΔCAPH2 mutant cells, at a resolution of 250 kb. Green lines represent inter-chromosomal interactions. To improve clarity, only randomly sampled 1% of the top 50% strongest interactions (edges) are shown. (c-d) Scatter plots showing the correlation between C4_E_ (L4E) and ΔC4 (D4) in two chromosomal systems: (c) chr17-chr22 system, and (d) chr19-chr22 system.

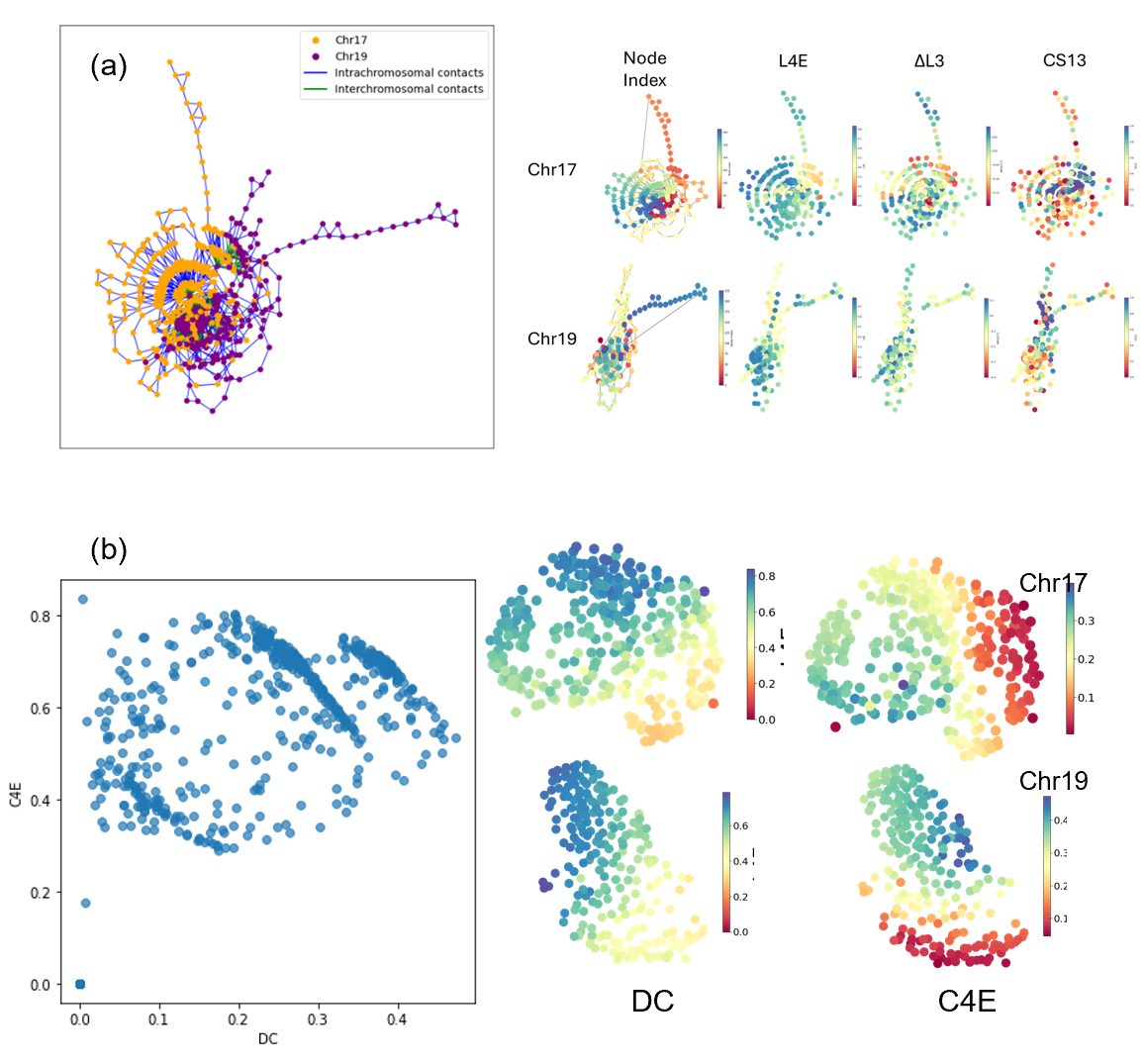

Figure S3:(a) Network display using the top 1% of links for a comparison to the random 1% of the top 50% used in Figure 3a. (b) The relationship between DC and C4_E_ for the chr17-chr19 cis contact interaction.

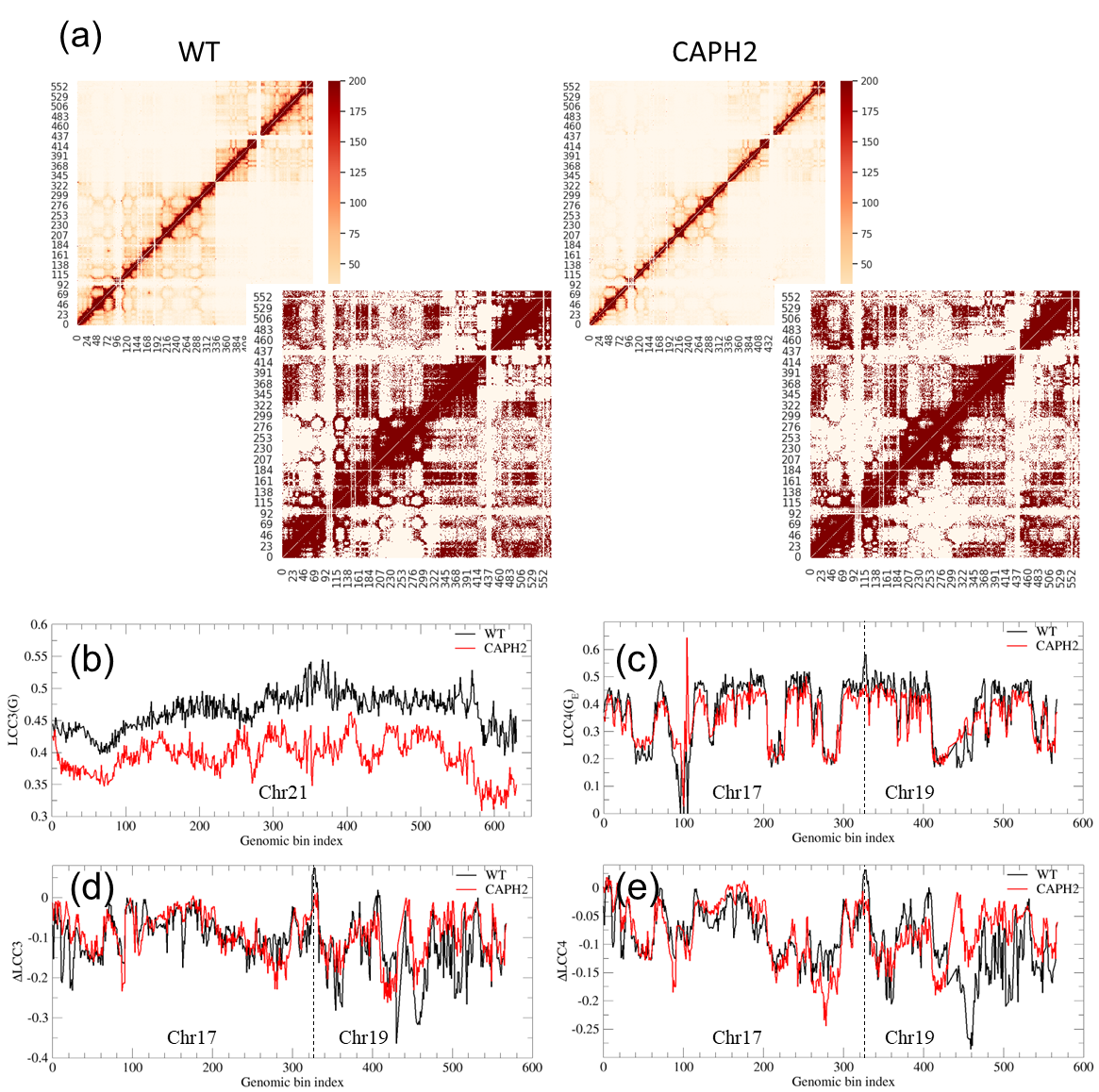

Figure S4: The comparison of chr17-19 interchromosomal structure networks between HAP1 wild type (WT) and a mutant (ΔCAPH2). Bin indices 0-324 correspond to chromosome17 (0-81,195,210 kb), and indices 325-561 correspond to chromosome 19 (0-59,128,983 kb), at a resolution of 250 kb. (a) Heatmap from Hi-C data and the associated contact network with 50% non-zero value cutoff. (b) Comparison of LCC3 (C3) values in the intrachromosomal contact network of chromosome 21, at a resolution of 50 kb. (c-e) Comparison of LCC4 (C4), ΔC3, and ΔC4 values in the interchromosomal contact networks of chr17-chr19 at 250-kb resolution.

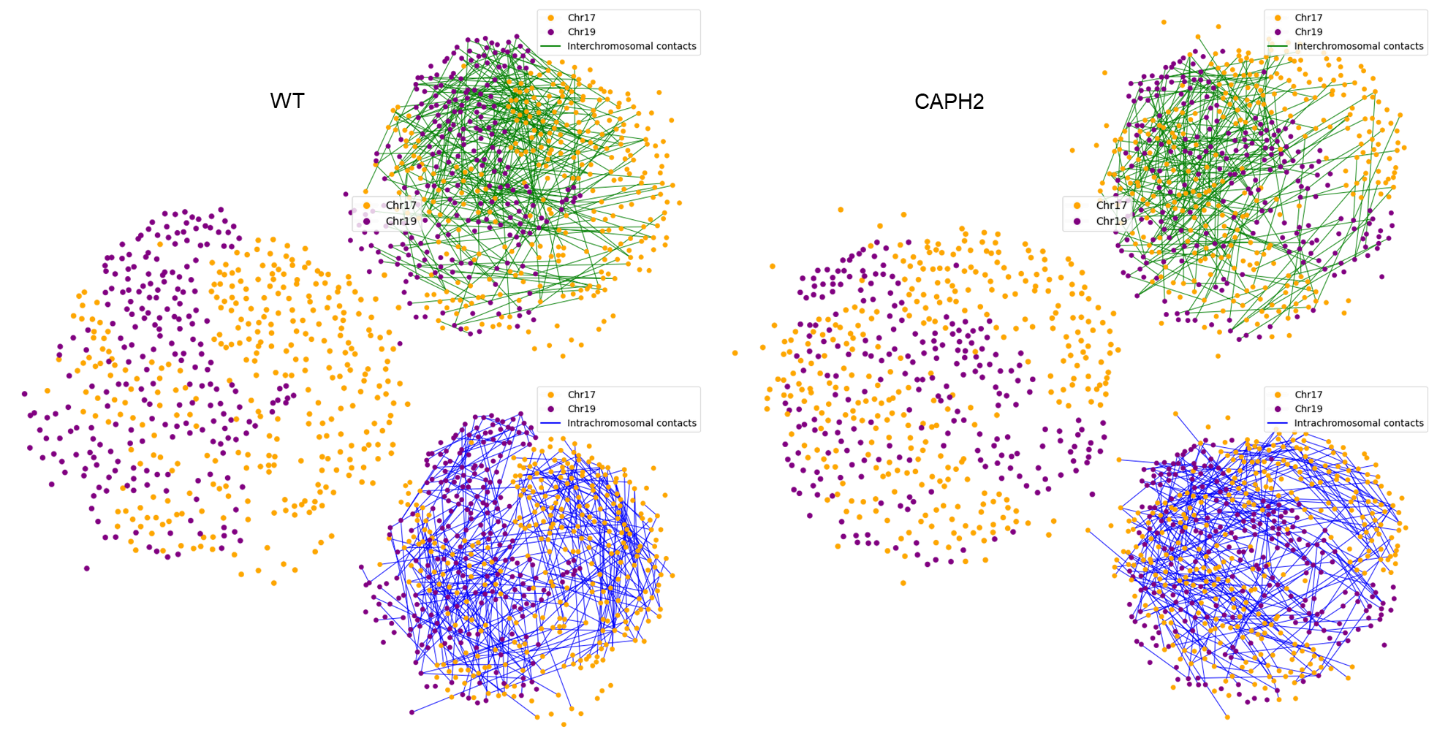

Figure S5: Force-directed layout of the chr17-chr19 contact network for HAP1 wild-type and ΔCAPH2 mutant cells, at 250-kb resolution. Green lines represent interchromosomal interactions, and blue lines represent intrachromosomal interactions. To improve clarity, only a randomly sampled 1% of the top 50% strongest interactions (edges) are shown.
